# Cell position-based evaluation of mechanical features of cells in multicellular systems

**DOI:** 10.1101/2024.11.10.622875

**Authors:** Hiroshi Koyama, Atsushi M. Ito, Hisashi Okumura, Tetsuhisa Otani, Kazuyuki Nakamura, Toshihiko Fujimori

## Abstract

Measurement of mechanical forces of cell–cell interactions is important for studying the emergence of diverse three-dimensional morphologies of multicellular organisms. We previously reported an image-based statistical method for inferring effective pairwise forces of cell–cell interactions (i.e., attractive/repulsive forces), where a cell particle model was fitted to cell tracking data acquired by live imaging. However, because the particle model is a coarse-grained model, it remains unclear how the pairwise forces relates to sub-cellular mechanical components including cell–cell adhesive forces. Here we applied our inference method to cell tracking data generated by vertex models that assumed sub-cellular components. Through this approach, we investigated the relationship between the effective pairwise forces and various sub-cellular components: cell–cell adhesion forces, cell surface tensions, cell–extracellular matrix (ECM) adhesion, traction forces between cells and ECM, cell growth, etc. We found that the cell–cell adhesion forces were attractive, and both the cell surface tensions and cell–ECM adhesive forces were repulsive, etc. These results indicate that sub-cellular mechanical components can contribute to the effective attractive/repulsive forces of cell–cell interactions. This comprehensive analysis provides theoretical bases for linking the pairwise forces to the sub-cellular mechanical components: this showcase is useful for speculating the sub-cellular mechanical components from the information of cell positions, and for interpreting simulation results based on particle models.

## 1. Introduction

In multicellular living systems, mechanical features of cells are critical parameters for morphogenesis. For instance, polygonal epithelial cells show a tightly packed cell configuration, where both cell–cell adhesive forces and cell surface tensions are major components for regulating cell shapes, fluidity of the tissues, and tissue deformation (Bi et al., 2015; Landsberg et al., 2009; Lecuit et al., 2011). To address the roles of the mechanical components of cells in the behaviors of whole systems, several frameworks for simulating multicellular systems have been developed such as vertex models, cell particle models, cellular Potts models, phase field models, etc. (Merkel and Manning, 2017; Nonomura, 2012).

To perform simulations, mechanical parameters are required that are measured by using techniques such as atomic force microscopies (Campàs, 2016; Sugimura et al., 2016) or by using image-based force inference methods (Brodland et al., 2010; Ishihara and Sugimura, 2012; Koyama et al., 2023). The latter techniques enable us to acquire spatio-temporal information of parameter values. For instance, by fitting a vertex model to microscopic images of labeled cell membranes, cell–cell junction tensions were inferred (Ishihara and Sugimura, 2012). In the case of three-dimensional (3D) tissues, imaging cell membranes with subsequent segmentation are challenging, while we previously reported a particle model-based force inference method where only cell nuclear positions were used as described in detail later (Koyama et al., 2023). In other words, this inference method is based on nuclear/cellular positions with their temporal evolution. However, because a particle model is a coarse-grained model, it is not clear how the forces in the particle model relate to the sub-cellular mechanical parameters including the cell–cell adhesion forces, the cell surface tensions, etc.

The cell particle models, where a cell is described as a point particle, have been widely applied to various cells including blastomeres, cancer cells, mesenchymal cells, and epithelial cells (Drasdo et al., 1995; Forgacs and Newman, 2005; Guillot and Lecuit, 2013; Maître et al., 2012; Merkel and Manning, 2017; Steinberg, 2007; Yong et al., 2015). The particle models typically assume both attractive and repulsive forces of cell–cell interactions: i.e., mechanical potential of pairwise cell–cell interactions. Because of their simplicity, low coding cost, and low computational load, the particle models are advantageous for simulating larger three-dimensional systems (Delile et al., 2017; Nissen et al., 2018). However, the origin(s) of the attractive/repulsive forces has not been correctly clarified yet. For instance, questions to be addressed include whether the attractive behavior of cell–cell interactions is conferred from the cell–cell adhesion forces, etc. This question is important to correctly interpret results of simulations based on the particle models in the context of the sub-cellular mechanical features.

In the case of isolated two cell systems (Fig. 1A-B), the cell–cell adhesion forces should shorten the cell–cell distance, which can be described as an effective attractive force, whereas both the cell surface tensions and excluded volume effect of the cells can be described as an effective repulsive force, as described in text books and in some articles (Forgacs and Newman, 2005; Merkel and Manning, 2017). Then, the summation of these different forces may lead to distance-dependent force curves (Fig. 1C). We previously reported that the profile of the distance–force curves is critical for morphogenesis during early embryogenesis in mice and *C. elegan*s (Koyama et al., 2024, 2023). However, it remains unclear whether the above relationships considered in the isolated two cell systems are applicable to multicellular systems (Fig. 1D), because the cells in multicellular systems can exhibit shapes far from spheres and so on.

**Figure 1:**
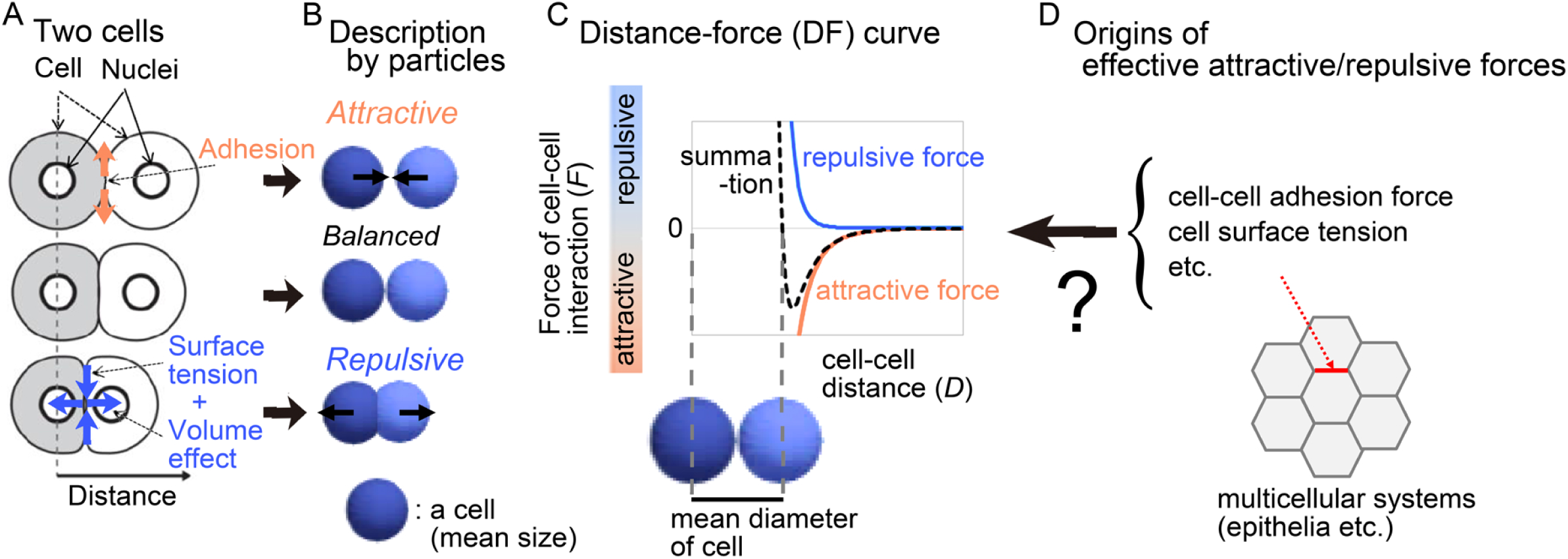
Attractive/repulsive forces of cell–cell interactions A. Sub-cellular forces contributing to cell–cell interactions are exemplified in the case of an isolated two cell system. B. The sub-cellular forces are approximated as attractive/repulsive forces in particle models. C. Both the repulsive and attractive forces are provided as cell–cell distance-dependent functions, and the summation (black broken line) is defined as the distance–force curve of the two particles. The relationship between the curve and the mean diameters of the cells is shown. D. The sub-cellular mechanical components in multicellular systems include the cell–cell adhesion forces, the cell surface tensions, etc. They can be the origin of the effective attractive/repulsive forces.

In the present study, we aimed to extend the cell position-based force inference framework. By using our image-based method for inferring effective distance–force curves (Koyama et al., 2023), we theoretically evaluated the relationships between the effective distance–force curves and the sub-cellular mechanical components. In other words, we generated synthetic data by simulating vertex models where both the cell–cell adhesion forces and the cell surface tensions were explicitly assumed, and then, by applying our inference method to the synthetic data, we evaluated the relationship between the distance–force curves and the sub-cellular mechanical components (Fig. 1D). In addition to these components, we also consider following sub-cellular components: the area elasticity (conservation of cell volume/area), cell growth (cell volume increase), adhesion forces between cells and extracellular matrix (ECM), and traction forces between cells and ECM. This comprehensive analysis provides the information whether each sub-cellular component effectively acts as the attractive or repulsive forces, and also provide relationships between the particle models and other models including the vertex models.

## 2. Theory

### 2.1. Method for inferring attractive/repulsive forces of cell–cell interactions

To infer the effective attractive and repulsive forces of cell–cell interactions in multicellular systems, we previously developed an image-based inference method (Koyama et al., 2023). Several studies have been adopting image-based strategies for inferring cellular mechanical forces and properties (Brodland et al., 2010; Ishihara and Sugimura, 2012). In the case of our method, a particle model was fitted to the temporal evolution of cell positions obtained from live microscopic imaging, which enabled us to infer the attractive/repulsive forces between each cell–cell interaction with each time frame as explained below. By analyzing the inferred attractive/repulsive forces, we reported that the effective attractive/repulsive forces can be actually approximated as a function dependent on the cell–cell distances (i.e., distance–force curves similar to Fig. 1C) in the blastomeres of the mouse and of *C. elegans* embryos (Koyama et al., 2024, 2023).

Briefly, in our method, we minimized a following cost function; *G* = *G_xyz_* + *G_F_*, where *G_xyz_* is the difference in *xyz*-coordinates of each particle between *in vivo* data and particle simulations, and *G*_*F*_0__ is the additional cost function for cell–cell distance-dependent weight (Koyama et al., 2023). *G_xyz_* was defined so as to contain the *xyz*- coordinates throughout all time frames:

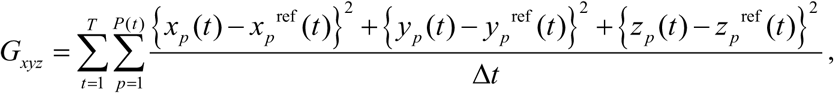

where *p* is the particle ID, *P*(*t*) is the total number of the particles at *t*, *T* is the total time frames, and Δ*t* is the time interval. *x_p_*, *y_p_*, and *z_p_* are the *xyz*-coordinates of the *p*th particles in the particle simulations. *x_p_*^ref^, *y_p_*^ref^, and *z_p_*^ref^ are the *xyz*-coordinates of the *p*th particles in the *in vivo* cells. The distance-dependent weight *G*_*F*_0__ means that the forces of the cell–cell interactions should be gradually decreased as the distances of the cell–cell interactions are increased, and ultimately, the forces become zero at the long distances. *G*_*F*_0__ was defined as follows: 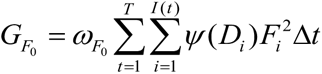, where *F_i_* is the force value of *i*th cell–cell interactions, *I*(*t*) is the total number of the cell–cell interactions at *t*, *D_i_* is the cell–cell distance of the *i*th interaction, and *ω_F_*_0_ is a coefficient for determining the relative contribution of the second term to *G*. *ψ*(*D_i_*) is a distance-dependent weight that exponentially increases, leading to that the value of *F_i_* approaches zero around larger *D_i_* (Koyama et al., 2023). We numerically solved this minimization problem by using the conjugate gradient method. Our inference method was previously validated by applying this method to simulation data generated under pregiven forces (Koyama et al., 2023). We had also confirmed that a unique solution was obtained by our minimization method (Koyama et al., 2023). The code is the same as that in Koyama et al. 2023 (https://doi.org/10.5281/zenodo.7427050).

### 2.2. Particle model to be fitted

In our particle model, attractive and repulsive forces for pairwise cell–cell interactions are assumed. Due to the low Reynolds number in the case of cellular-level phenomena, the inertial force is negligible (Fletcher et al., 2014; Forgacs and Newman, 2005; He et al., 2014; Li et al., 2017; Schötz et al., 2008). In our present study, we assumed that cells were put on substrates, and that frictional forces between the cells and the substrates were exerted, by which the motion of the particles was overdamped. In the case that the substrates are fixed (i.e., non-moving), the equation of a particle motion may be written as follows;

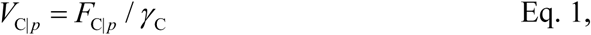

Where *V*_C|_*_p_* is the velocity of the *p*th particle, *F*_C|_*_p_* is the summation of the cell–cell interaction forces exerted on the *p*th particle, and *γ*_C_ is the coefficient of the viscous drag and frictional forces provided from the substrates. This assumption has been used in both particle-based models and vertex models (Honda et al., 2008; Li et al., 2017; Nissen et al., 2017; Suzuki et al., 2017; Szabó et al., 2006).

During this inference, we assumed the simplest situation, where *γ*_C_ is constant (=1.0). This is the same setting as that in our previous work (Koyama et al., 2023). The dimension of the particle velocity *V*_C|_*_p_* was set to be μm/min for convenience, which is typical dimension in living cells. Because we did not measure the value of *γ*_C_ in living tissues, we cannot determine the value of the effective force *F*_C|_*_p_* (= *γ*_C_*V*_C|*p*_) with a specific dimension. However, under the above assumption, 1 arbitrary unit (A.U.) of *F* coincides with a force which can move a particle with a velocity of 1 μm/min in tissues.

### 2.3. Vertex model simulation as synthetic data

As a well-defined system, we adopted the vertex model for describing epithelial cells (Fig, 2A, “Model”). This model is the standard framework for simulating epithelial cells which typically exhibit polygonal shapes, where some sub-cellular mechanical components such as cell–cell adhesion forces and cell surface tensions are assumed (Fletcher et al., 2014; Okuda et al., 2015). By using the vertex model, we performed simulations, and we considered the simulation outcomes as synthetic data (Fig. 2A, “Simulation”). In other words, we applied our inference method to cell tracking data generated by the simulations of the vertex model, and then obtained effective distance–force curves (Fig. 2A, “DF curve”).

**Figure 2:**
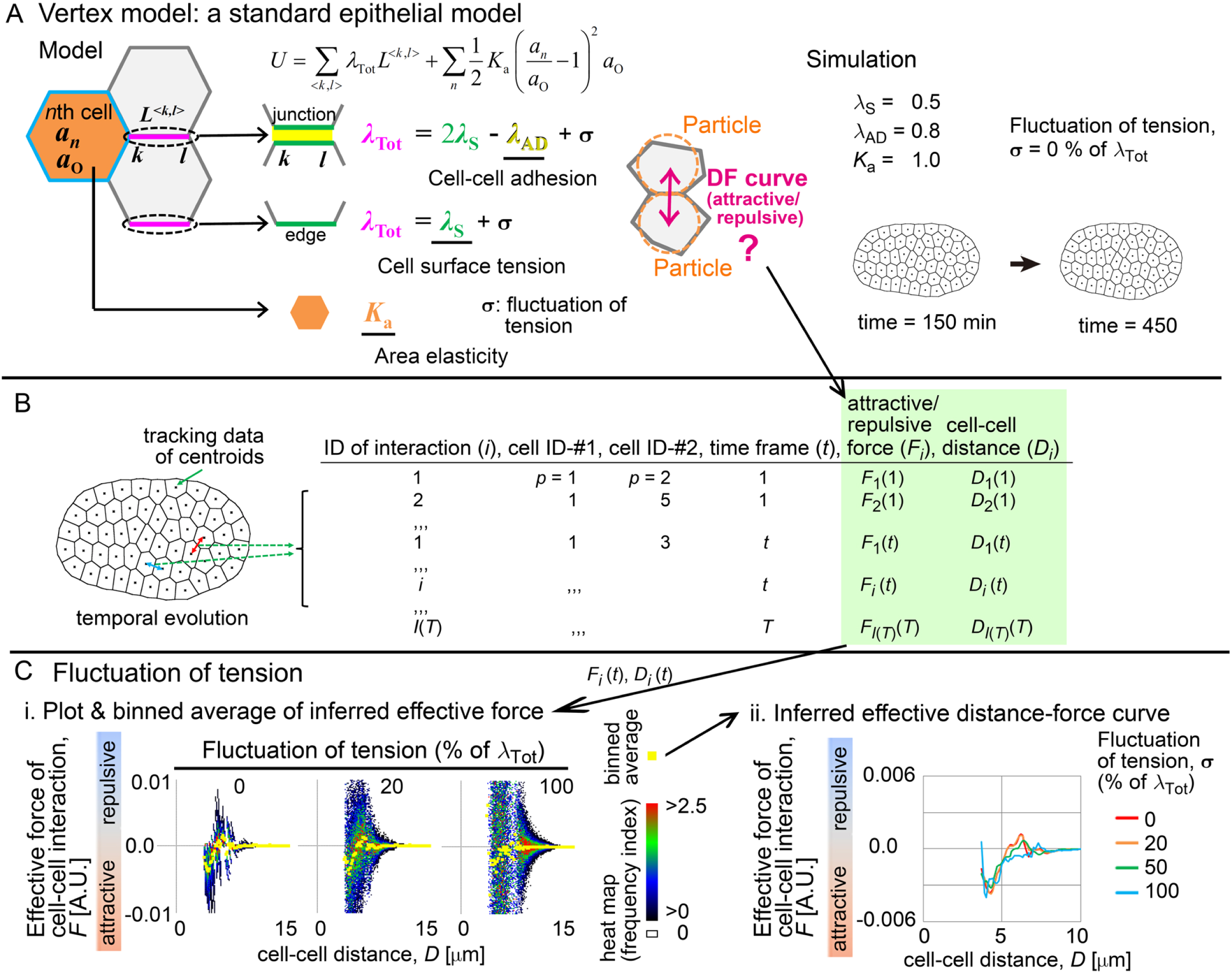
Inference of effective attractive/repulsive of cell–cell interaction in vertex model A. Vertex model. *λ*_Tot_ is the line tension exerted on the cell–cell junctions or on the cellular edges along the outer boundary of the cell cluster. The potential energy on a junction or edge is *λ*_Tot_*L*^<^*^k^*^,*l*>^, where *L* is the length of the junction or the edge composed of the *k*th and *l*th vertices. The potential energy for a cell provided by the area elasticity is (1/ 2)*K_a_* (*a_n_*/ *a*_o_−1)^2^ *a*_o_, where *a_n_* is the area of the *n*th cell and *a*_o_ is the natural area. An example of the simulations is shown with the parameter values (right panel). In addition to the polygonal cell contours, the cell centroids are shown as dots (See B). B. Structure of the data obtained by our inference method. *F_i_*(*t*) is the inferred force of the *i*th cell–cell interaction at *t*, where the distance is *D_i_*(*t*). C. Inferred effective distance–force (DF) curves under various values of the fluctuations of the line tensions, *σ*. i) The inferred forces are plotted against the cell–cell distances, and the values of the binned averages of the forces are shown in yellow. The width of the bins for the distances (*D*) is 0.27 μm. The procedure is detailed in the Supplementary information. ii) The inferred DF curves were shown, which were obtained by linearly connecting the plots of the binned averages in (i).

The vertex model used in this study is essentially the same as that in our previous studies (Koyama et al., 2022; Koyama and Fujimori, 2020). The potential energy in the whole system is defined as follows,

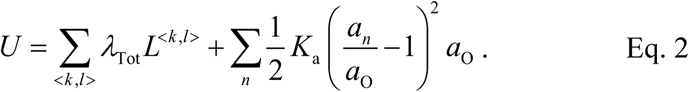

*λ*_Tot_ is the line tension exerted on cell–cell junctions or on cell edges of outer boundary (Fig. 2A, “Model”). The line tension is provided from both the cell–cell adhesion force (*λ*_AD_) and the cell surface tension (*λ*_S_); the former has a negative effect on the line tension, and the latter does a positive (Fig. 2A; *λ*_Tot_ = 2*λ*_S_ − *λ*_AD_). The potential energy on a junction or edge is *λ*_Tot_*L*^<^*^k^*^,*l*>^, where *L* is the length of the junction or edge composed of the *k*th and *l*th vertices. The potential energy for a cell provided by the area elasticity is (1/ 2)*K_a_*(*a_n_*/ *a*_o_−1)^2^ *a*_o_, where *a_n_* is the area of the *n*th cell and *a*_o_ is the natural area. The force (*F*_V|*h*_) exerted on an *h*th vertex is calculated as follows: *F*_V|*h*_ = −∇*U*, where ∇ is the nabla vector differential operator at each vertex. Similar to the particle model, we assumed that the cells were put on non-moving substrates where frictional forces were exerted. Therefore, the motion of each vertex is described as follows: *V*_V|*h*_ = *F*_V|*h*_ / *γ*_V_, *V*_V|*h*_ is the velocity of the *h*th vertex, and *γ*_V_ is the coefficient of the friction of a vertex. Similar to the case in the particle model, *γ*_V_ was set to 1.0, and the dimension of particle velocity *V*_V|*h*_ was set to be μm/min. The code is the same as that given in our previous work (Koyama and Fujimori, 2020).

The fluctuation of the line tensions (*σ*) was assumed as follows. Cellular edges to be activated were selected according to *p*_ED_ which is the probability of activation per unit of time. On the cellular edges selected, *σ* was added to the line tensions, and the duration time of this activated state is *τ* as defined in the Supplementary information.

## 3. Results

### 3.1. Inference of attractive/repulsive forces in vertex model

To evaluate possible relationships between the attractive/repulsive forces and the cell– cell adhesion forces/the cell surface tensions, we applied our inference method to synthetic cell tracking data generated by our vertex model. We performed simulations containing several tens of cells (Fig. 2A, “Simulation”). Then, by tracking the centroids of each polygonal cell, we generated cell tracking data (Fig. 2B). By applying our inference method to the cell tracking data, we obtained the attractive/repulsive forces for each cell–cell interaction at each time frame (Fig. 2B, “*F_i_*(*t*)”), and we simultaneously acquired the distances for each cell–cell interaction (Fig. 2B, “*D_i_*(*t*)”). These paired data (i.e., *F_i_*(*t*) and *D_i_*(*t*)) were plotted on a graph space below.

Fig. 2C-i shows the plots of the inferred forces (*F_i_*(*t*)) against the cell–cell distances (*D_i_*(*t*)). In the absence of the fluctuations of tensions (0%), the distribution of the data points was not so diverged (Fig. 2C-i, heat maps), and, by calculating the values of binned averages, a DF curve was detected (Fig. 2C-i, yellow). The binned averaged line in Fig. 2C-i is shown in Fig. 2C-ii (red line), which exhibited a typical profile with a peak of the attractive force. In the presence of the fluctuations, the distributions became wider (Fig. 2C-i, heat maps), but the DF curves were detected whose profiles were equivalent to that without the fluctuations (Fig. 2C-ii). In conclusion, the effective DF curves are detectable in multicellular epithelial cells.

### 3.2. Influence of cell–cell adhesion forces/cell surface tensions on DF curves

Next, we examined whether the microscopic features of the epithelial cells were reflected in the effective DF curves. We performed simulations under various values of the cell– cell adhesion forces (Fig. 3A-i); the cells showed highly deformed cell shapes or regular hexagonal shapes under the stronger or weaker cell–cell adhesion forces, respectively. The inferred DF curves are shown in Fig. 3A-ii, whose profiles are quantitatively evaluated in Fig. 3A-iii. The absolute values of the maximum attractive forces were positively correlated with the cell–cell adhesion forces (Fig. 3A-iii). These results are consistent with the case in the isolated two cell systems.

**Figure 3:**
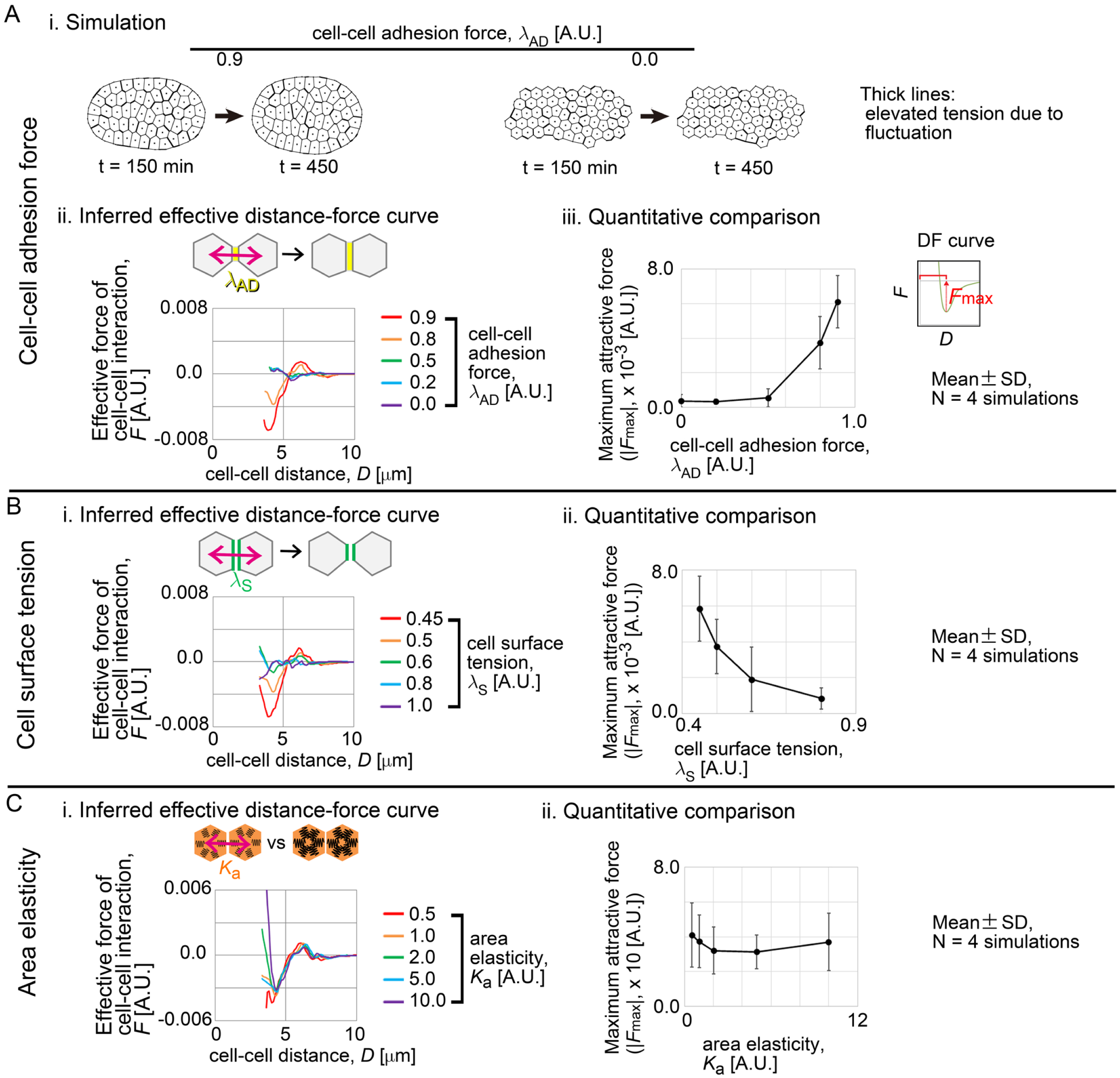
Effective attractive/repulsive forces in vertex model A. Inferred DF curves under various values of the cell–cell adhesive forces, *λ*_AD_. i) Simulation outcomes are shown where the cell cluster tends to form a circular colony under the increased values of the adhesive forces (*λ*_AD_ = 0.9). ii) Inferred DF curves. iii) Quantitative comparison of the DF curves. The maximum attractive force (*F*_max_) is defined. B. Inferred DF curves under various values of the cell surface tensions, *λ*_S_. i) Inferred DF curves. ii) Quantitative comparison of the DF curves. C. Inferred DF curves under various values of the area elasticities, *K*_a_. i) Inferred DF curves. ii) Quantitative comparison of the DF curves. Related Figure: Figure S1 (inferred DF curves in the presence of the potential derived from the cell perimeters).

We also evaluated the effect of the cell surface tensions (Fig. 3B), and successfully detected DF curves (Fig. 3B-i). The maximum attractive forces were negatively correlated with the cell surface tensions. Therefore, the effect of the cell surface tensions was opposite to that of the cell–cell adhesion forces (Fig. 3B-i vs. 3B-ii). If the surface tensions act as the repulsive forces, they can be emerged as the reduction of the attractive forces in the DF curves, because the DF curves are the summation of the repulsive and attractive forces (Fig. 1C). Together with the results in the case of the cell– cell adhesion forces, these results indicate that the effects of the microscopic features of the cells are conserved among the multicellular epithelial cells and the isolated two cell systems.

Finally, we evaluated the effect of the area elasticity. Under large values of the area elasticity, one expectation is that the slope of the repulsive region in the DF curve becomes steeper due to its elasticity. The inferred effective DF curves and their quantitative evaluations are shown in Fig. 3C-i and 3C-ii. The typical profiles of the DF curves such as the peak of the attractive forces were clearly detected, but the maximum values were not significantly affected by the area elasticity (Fig. 3C-ii). On the other hand, although the profiles of the repulsive regions were not obtained in all conditions, the slope under the higher value of the area elasticity (= 10.0) was steepest (Fig. 3C-i, purple line). Therefore, the effect of the area elasticity is neutral on the attractive forces in the DF curves, and, because the slope of the repulsive region is related to the stiffness of the cores of the cells, the area elasticity may be correlated with the stiffness of the cores.

In summary, the cell position-based method for evaluating the mechanical properties of multicellular epithelial cells revealed that the cell–cell adhesion forces and the cell surface tensions showed attractive and repulsive effects, respectively. In addition, we also tested whether these relationships were conserved in other vertex models (i.e., model dependency). We used another typical vertex model that assumed a potential based on the perimeters of the cells, which increased the cell–cell junction tensions (Fletcher et al., 2014). The definition of the potential is described in the Supplementary information. If the coefficient of this potential was not so large, the relationships were conserved (Fig. S1).

### 3.3. Influence of cell-ECM adhesion on DF curve

In real tissues, there are various types of non-cellular external factors which affect cell behaviors; extracellular matrix (ECM), egg shells, liquid cavity, etc (Honda et al., 2008; Kajita et al., 2003). These factors may influence inferred effective DF curves. This is analogous to the atomic, molecular, and colloidal sciences. For instance, external pressures increase the density of particulate objects, which is reflected in the radial distribution function linked to the effective distance–potential curves (Atkins et al., 2018). Another example is solvents, where hydration of particles by the solvents yields energy barriers in the effective distance–potential curves (Israelachvili, 2011). Therefore, by measuring the effective distance–potential curves, one can predict, at least partially, what kind of external factors exist in systems of interest. In multicellular systems, however, it has not been clarified how external factors influence the effective distance–potential curves.

Here we examined the influence of ECM on the effective DF curves, that interacts with cells. Spreading of cells on ECM is regulated by the competition between the cell–ECM and the cell–cell adhesions; the increased forces of the cell–ECM adhesions lead to the increase in the areas of the cells (Fig. 4-i) (Pérez-González et al., 2019; Ryan et al., 2001; Yong et al., 2015). Because the increase in each cell area pushes the surrounding cells, we expected that the cell–ECM adhesion forces would show a repulsive effect on the effective cell–cell interactions (Fig. 4-i, “Repulsive force?”). Simultaneously, the increased cell areas lead to the increases in the distances between the cells, which should be appeared as a rightward shift of the effective DF curves along the cell–cell distances (Fig. 4-i, “Shift of DF curve?”).

**Figure 4:**
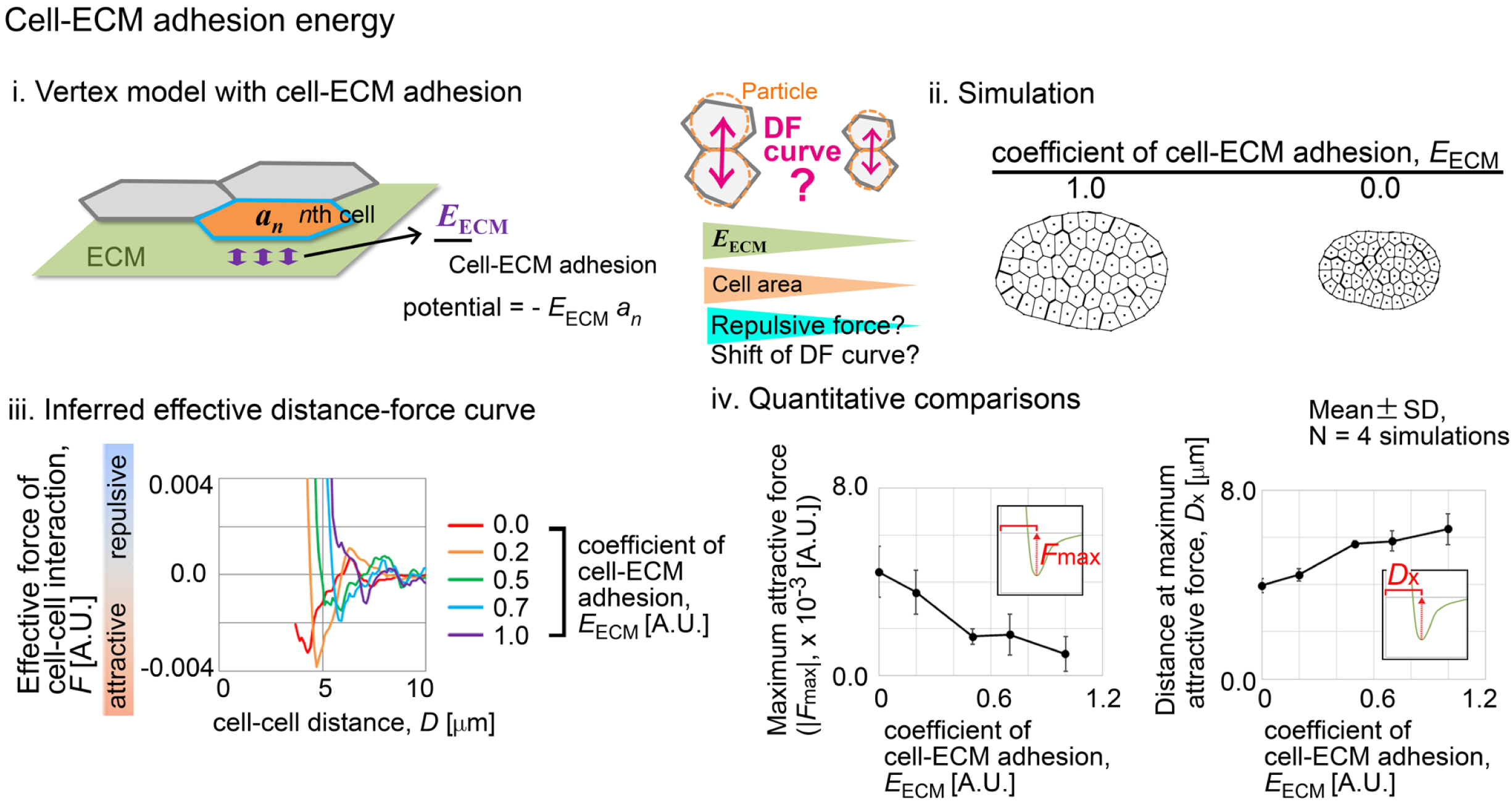
Effective attractive/repulsive forces in the presence of cell-ECM adhesion The cell–ECM adhesion potential energy was assumed in the vertex model used in Fig. 2-3. i) The potential for a cell is −*E*_ECM_ *a_n_*, where *E*_ECM_ is the coefficient of the potential. ii) Simulation outcomes are exemplified where the cell areas are increased under the larger values of *E*_ECM_. The left and right panels are shown at the same scale. iii) Inferred DF curves. iv) Quantitative comparisons of the DF curves. The insets illustrate DF curves, where *F*_max_ and *D*_x_ are defined. Related Figure: Figure S2 (The effect of the cell surface tensions on the distances at the maximum attractive forces is shown. See Fig. 4-iv and Fig. S2.).

We modeled the cell–ECM adhesion in the vertex model used in Fig. 2. The adhesion between cells and ECM is mediated by focal adhesions, etc (Goodwin et al., 2016; Pérez-González et al., 2019; Yamaguchi et al., 2022). The distributions of these adhesive structures seem to be different among tissues and conditions: some tissues show distributions along their peripheries, while others show almost unbiased distributions. Here we assumed the simplest case where the structures are uniformly distributed. We defined the potential energy of the adhesion as −*E*_ECM_ *a_n_* for each cell, where *E*_ECM_ is the coefficient of the energy and *a_n_* is the cell area as previously defined.

Under the increased cell–ECM adhesion forces, the cell areas were increased (Fig. 4-ii, *E*_ECM_ = 1.0 vs. = 0.0). Fig. 4-iii shows the inferred effective DF curves, and Fig. 4-iv is the quantitative evaluation of the curves. As we expected, the maximum attractive forces were negatively correlated with the cell–ECM adhesive forces (Fig. 4-iv, left panel), and the distances at the maximum attractive forces were increased by the cell–ECM adhesive forces (Fig. 4-iv, right panel). These results suggest that the cell–ECM adhesive forces are reflected in the reduction of the attractive forces in the effective DF curves.

In addition, according to Fig. 4 and Fig. 3B, both the cell–ECM adhesive forces and the cell surface tensions showed the repulsive effects. However, the latter did not show an increase in the distances at the maximum attractive forces (Fig. S2), suggesting that we can distinguish the two components each other.

### 3.4. Influence of cell-ECM traction forces on DF curve

Next, we focused on traction forces between cells and ECM, which confer self-migratory activities of the cells (Aoki et al., 2017; Garcia et al., 2015; Szabó et al., 2006). We implemented the traction forces in the particle model instead of the vertex model, because there is no general way to implement self-migratory activities in the vertex models. First, we set the traction forces to show random directions (Fig. 5A-i, right panel), and to persist for a given period of time (Fig. 5A-ii, “with persistency = 50.0min”). The cell–cell interaction forces were provided by the Lennard-Jones (LJ) potential (Fig. 5A-i, “LJ”). The value of the traction forces is shown as a relative value to the maximum attractive force of the LJ potential (Fig. 5A-ii, “1,000% of LJ”). We assumed that the friction between the cells and the ECM, and performed two-dimensional simulations with the friction between the cells and the ECM.

**Figure 5:**
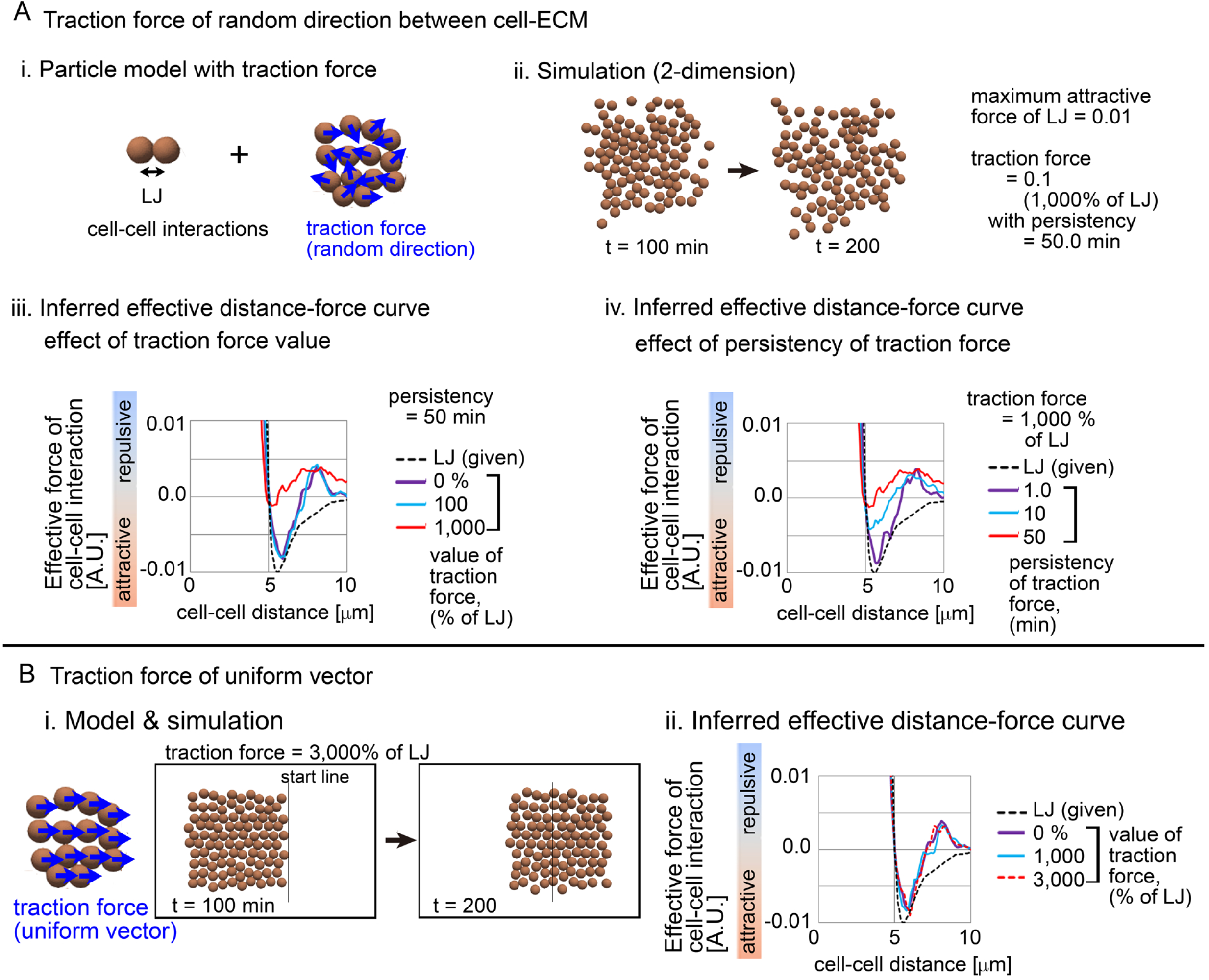
Effective attractive/repulsive forces in the presence of cell-ECM traction forces A. The traction forces between cells and ECM were assumed in the particle model under two-dimensional conditions. i) The cell–cell interaction forces were provided by the LJ potential. The directions of the traction forces were assumed to be random. ii) An example of the simulations is shown. iii and iv) Inferred DF curves under various values of the traction forces and under various values of the persistency, respectively. The distance– force curve provided by the LJ potential is also shown (broken black lines). B. The traction forces were assumed to have the same vector (i.e., the same value and the same direction). i) An example of the simulations is shown. Due to the same vector, the particles migrated unidirectionally. ii) Inferred DF curves under various values of the traction forces. The distance–force curve provided by the LJ potential is also shown. Related Figures: Figure S3 (inference in the self-migratory MDCK cells on substrates)

Fig. 5A-iii shows the inferred effective DF curves. In the absence of the traction forces (0%), the inferred DF curve showed a profile similar to the given LJ potential (Fig. 5A-iii, purple line), though an inconsistency was observed at the distant region (distance = ∼7μm). In the case that the value of the traction forces was equivalent to the attractive force of the LJ potential (100%; light blue line), the inferred curve was still consistent with that without the traction forces, whereas the inferred curve became highly repulsive under a larger traction force (1,000%; red line).

Fig. 5-iv shows the effect of the persistency of the traction forces. Even under the large traction forces (1,000%), if the persistency was short (1.0min; purple line), the inferred DF curve was consistent with the given LJ potential. In conclusion, the traction forces can significantly modify the effective DF curves under the condition that both the force value and the persistency are large. Otherwise, the traction forces do not affect the profiles of the DF curves. We also found such repulsive profiles of the DF curves in Madin-Darby canine kidney (MDCK) cells on substrates, which show highly self-migratory activities (Fig. S3A).

### 3.5. Influence of cell growth on DF curve

Cell growth (i.e., increase in cell volume during the cell proliferative phase) is an important parameter for homeostasis in multicellular systems: cell growth increases pressure in multicellular systems, leading to contact inhibition of growth, delamination of epithelial cells, and apoptosis (Marinari et al., 2012; Podewitz et al., 2015; Zimmermann et al., 2016). Under the increased pressure due to cell growth, the cells push/repel each other. In other words, the intrinsic attractive forces between the cells may be effectively cancelled due to the cell growth. To test this hypothesis, we used a mathematical model considering the cell growth. A well-known cell growth model is shown in Fig. 6-i, where one cell is modeled as two particles and the growth is described by the repulsive force (coefficient *B*) between the two particles (Basan et al., 2011; Podewitz et al., 2015; Ranft et al., 2010). The repulsive force is defined as follows; *B* / (*D*_g_ + *C*)^2^, where *D*_g_ is the distance between the paired two particles, and *C* is constant. When the distance between the two particles reaches a critical value, cell division occurs. In our model, we set *C* = 5.6 and *D*_CR_=5.0, and implemented that particle–particle interactions between different cells were governed by the LJ potential that was previously shown in Fig. 5A-iii, “LJ”. The original model by Basan et al. was developed for describing three-dimensional cell aggregates where viscous frictions between the cells are assumed. However, we modified this model so that the cell–substrate friction was assumed in a manner similar to Fig 2-5, instead of the frictions between the cells.

**Figure 6:**
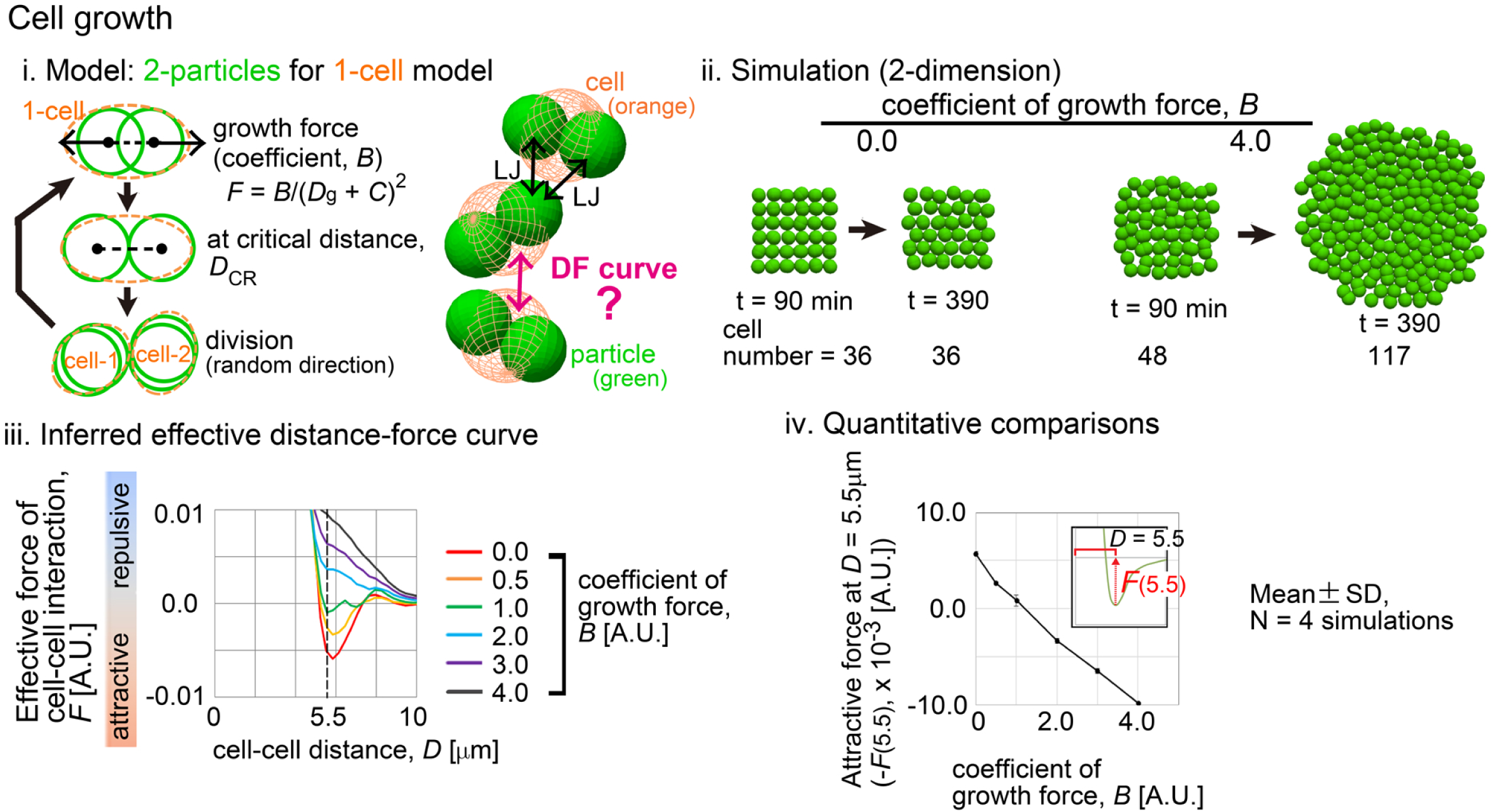
Effective attractive/repulsive forces in growing cells i) Model of cell growth. Orange, a cell; green, a particle. The growth was expressed as a repulsive force between the two particles as defined in the main text (*C* = 5.6 and *D*_CR_ = 5.0). The distance–force curve provided by the LJ potential was set to have the maximum attractive force = 0.005. When *B* = 0.5 and *D*_g_ = 4.4, the repulsive force given from the growth becomes 0.005, equivalent to the attractive force of the LJ. ii) Examples of the simulations under different values of *B*, where the particles but not the cells are visualized. iii) Inferred DF curves. iv) Quantitative comparison of the DF curves.

We performed simulations and, inferred effective DF curves between the cells (Fig. 6-i, “DF curve” between the orange cells). When the coefficient of the cell growth was set to be larger, the cell number was increased (Fig. 6-ii, *B* = 4.0 vs. 0.0). Fig. 6-iii shows the inferred effective DF curves, and Fig. 6-iv shows their quantitative comparison. The maximum attractive forces were negatively correlated with the growth forces, and under the larger growth forces, the attractive components were absolutely diminished (Fig. 6-iv). These results indicate that the cell growth effectively decreases the attractive forces of the cell–cell interactions.

### 3.6. DF curve in vertex model under steady state

In the above cases (Fig. 2-6), we assumed non-steady states: the systems are undergoing morphological changes. Such non-steady states should be typical during morphogenesis of multicellular systems. On the other hand, confluent epithelial monolayers would be under steady states, that may correspond to tissues in adult organs. Under the steady states, the motions of cells can be fluctuated, but the tissues do not experience morphological transitions; i.e., analogous to systems under thermodynamically equilibrium states. By using the vertex model, we then investigated the relationship between the effective DF curves and the cell–cell adhesion forces/the cell surface tensions under the steady states.

Fig. 7A shows a confluent epithelial monolayer modeled by the vertex model where the vertices continuously moved due to the fluctuations of the line tensions of the cell–cell junctions defined in Fig. 2. A whole view of the simulation is shown in Fig. S4.

**Figure 7:**
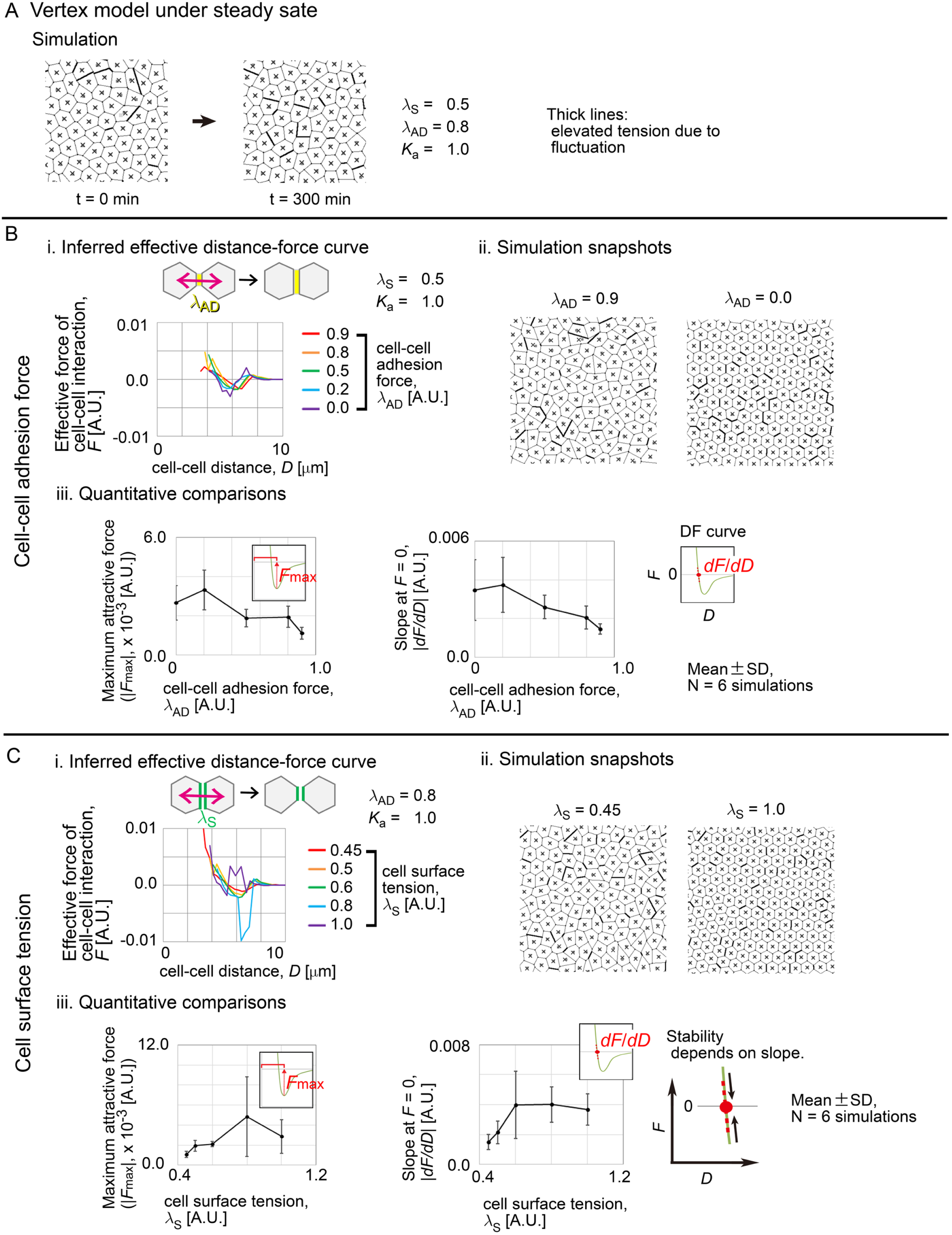
Effective attractive/repulsive forces under steady state A. Examples of vertex model simulations under steady state. The vertex model is the same as that in Fig. 2-3. The cell centroids are shown as crosses. The whole view of the simulation is shown in Fig. S4. B. Inferred DF curves under various values of the cell–cell adhesive forces, *λ*_AD_. i) Inferred DF curves. ii) Simulation outcomes. iii) Quantitative comparisons of the DF curves. The insets illustrate DF curves. C. Inferred DF curves under various values of the cell surface tensions, *λ*_S_. i) Inferred DF curves. ii) Simulation outcomes. iii) Quantitative comparisons of the DF curves. The slope (*dF*/*dD*) is related to the stability of the state with *D* at *F* = 0. Related Figures: Figure S4 (whole view of the simulation, and inferred DF curves under various values of the area elasticities, *K*_a_), Figure S5 (inferred DF curves in MDCK cells under the steady states)

#### 3.6.1. Influence of cell–cell adhesion forces/cell surface tensions on DF curve

Fig. 7B shows the inferred effective DF curves under the conditions with the different values of the cell–cell adhesion forces. In the all cases, the DF curves showed clear peaks of the attractive forces (Fig. 7B-i). In Fig. 7B-ii, the simulation snapshots show that the cells exhibited either highly deformed shapes or an array of regular hexagons under the stronger or weaker cell–cell adhesion forces (*λ*_AD_ = 0.9 vs. 0.0), respectively. The profiles of the DF curves were quantitatively evaluated (Fig. 7B-iii); unlike the case in Fig. 3A, there was a negative relationship between the cell–cell adhesion forces and the maximum attractive forces (Fig. 7B-iii, left panel). In addition, the slopes of the DF curves at the force = 0 —defined in Fig. 7B-iii, right panel, inset— showed a negative relationship to the cell–cell adhesion forces (Fig. 7B-iii, right panel). We will discuss these results later.

Fig. 7C shows the inferred effective DF curves under the conditions with the different values of the cell surface tensions. Clear DF curves were detected (Fig. 7C-i), except that the profiles were abruptly changed between the conditions of the weaker surface tensions and the conditions of the stronger surface tensions as follows (Fig. 7C- i). Under the three conditions with the weaker surface tensions (Fig. 7C-i, *λ*_S_ = 0.45, 0.5, and 0.6), the profiles of the three DF curves were almost similar, with gradual changes among them. Under the two conditions with the stronger surface tensions (Fig. 7C-i, *λ*_S_ = 0.8 and 1.0), the two DF curves showed an additional attractive component (*λ*_S_ = 0.8) or repulsive component (*λ*_S_ = 1.0) around the longer distances. Therefore, the profiles of the DF curves were abruptly changed, when the surface tensions were changed from [*λ*_S_ = 0.45, 0.5, 0.6] to [0.8, 1.0]. Under the stronger surface tensions, the cells showed an array of regular hexagons compared with the case under the weaker surface tensions (Fig. 7C-ii). The profiles of the DF curves were quantitatively evaluated (Fig. 7C-iii); unlike the case in Fig. 3C, there was a positive relationship between the surface tensions and the maximum attractive forces (Fig. 7C-iii, left panel). The slopes of the DF curves at the force = 0 showed positive relationship to the cell surface forces (Fig. 7C-iii, right panel).

We interpret the meaning of the slopes of the DF curves at the force = 0 as follows. In Fig. 7B-C, the maximum attractive forces show a negative or positive relationship to the cell–cell adhesion forces or the cell surface tensions, respectively. The increased maximum attractive forces (i.e., deeper valleys in the DF curves) are usually associated with steeper slopes of the DF curves, resulting in the steeper slopes at the distance at the force = 0 (Fig. 7B-iii and 7C-iii). Because the state of the force = 0 corresponds to a steady state, the slope at the force = 0 relates to the stability of the state; i.e., in the case of a system with the steeper negative slope, when the distance is slightly changed from the steady state, the force for restoring to the steady state becomes larger (Fig. 7C-iii, right panel, “Stability”). In other words, a system with a steeper slope strongly prefers to keep the cell–cell distances at a unique distance equal to that at the force = 0. This feature should lead to the formation of an array of regular hexagons as shown in Fig. 7B-ii and 7C-ii (*λ*_AD_ = 0 in B-ii or *λ*_S_ = 1.0 in C-ii). In conclusion, we think that a system that prefers a regular hexagonal array will show a steeper slope with a larger maximum attractive force.

In addition, we also evaluated the influence of the area elasticity (Fig. S4), and detected no significant changes in the inferred DF curves under the different values of the area elasticities. We also inferred DF curves in MDCK epithelial cells under a nearly stable state, and detected a clear DF curve with the attractive forces (Fig. S5).

#### 3.6.2. DF curve in solid-liquid phase transition

As mentioned above in Fig. 7C-i, we detected an abrupt change in the profiles of the inferred DF curves. It is possible that such an abrupt change corresponds to a certain phase transition. In the case of multicellular systems, solid-liquid phase transitions are well known, which are also reported in the vertex models (Bi et al., 2015; Mongera et al., 2018; Petridou et al., 2021). We evaluated the fluidity of the cells in our vertex model-based simulations. Fig. 8A shows the trajectories of the cell centroids under the conditions of the different values of the cell surface tensions. Under the smaller values of the surface tensions (*λ*_S_ = 0.45, 0.5), the cell centroids were substantially moved during the simulations, indicating a liquid-like behavior. On the other hand, under the larger values of the surface tensions (*λ*_S_ = 0.8, 1.0), the positions of the cell centroids were almost fixed, indicating a solid-like behavior. Fig. 8B shows the mean square displacement (MSD). The MSDs were continuously increased under the weaker surface tensions (*λ*_S_ = 0.45, 0.5, 0.6), whereas the MSDs reached plateaus under the stronger surface tensions (*λ*_S_ = 0.8, 1.0). Therefore, a solid-liquid transition occurs between *λ*_S_ = 0.6 and 0.8, and this critical point corresponds to the abrupt change in the profiles of the inferred DF curves. We concluded that the solid-liquid phase transition was appeared as the abrupt change in the inferred DF curves.

**Figure 8:**
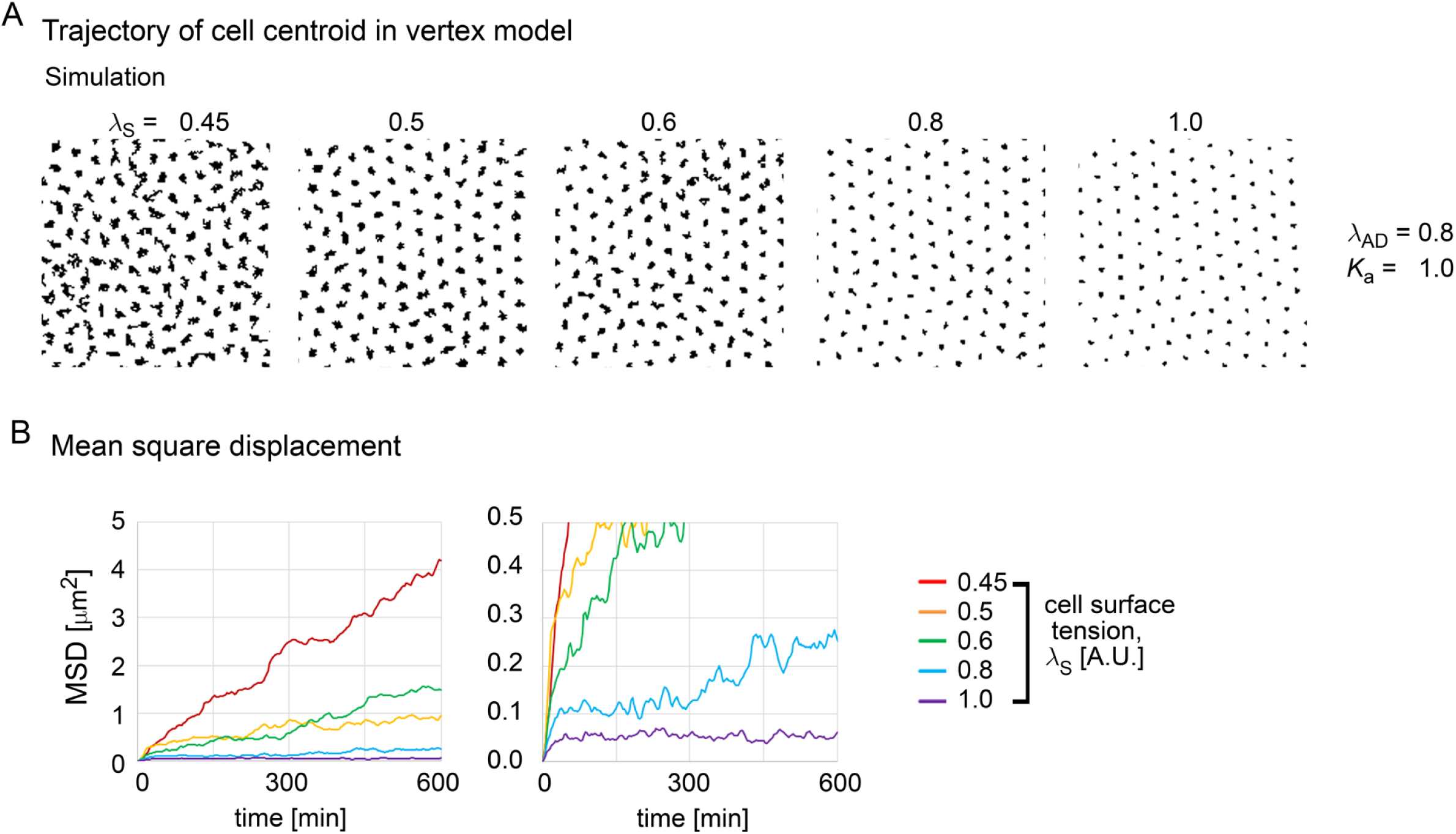
Solid-liquid phase transition in vertex model A. The trajectories of the cell centroids in the vertex model in Fig. 7 are shown under various values of the cell surface tensions. The sizes of each window correspond to those in the simulation snapshots in Fig. 7. B. The mean square displacements (MSD) are shown under various values of the cell surface tensions.

## 4. Discussion

In the present study, we investigated how the sub-cellular mechanical components relate to the effective attractive/repulsive forces of cell–cell interactions in multicellular systems. By applying our particle model-based inference method to the cell tracking data generated by some model frameworks, such as the vertex model and the 2-particel-for-1-cell model, we related the effective attractive/repulsive forces to the various sub-cellular mechanical components. Our comprehensive analyses are summarized in Table 1: the cell–cell adhesion forces effectively act as attractive forces, the cell surface tensions act as repulsive, the cell growth acts as repulsive, the cell–ECM adhesion forces act as repulsive, the stronger traction forces with long persistency act as repulsive. In Table 1, we also added the effect of egg-shells and liquid cavities as we previously reported (Koyama et al., 2024, 2023). Therefore, our results indicate that some sub-cellular mechanical components actually act as the effective pairwise forces of the cell–cell interactions. The quantitative differences in the profiles of the inferred DF curves are critical for morphogenesis (Koyama et al., 2024, 2023). When performing the cell position-based force inference method, this comprehensive description would be useful to speculate the origins of the attractive/repulsive forces.

**Table 1:** Effect of cellular and non-cellular factors on effective force of cell–cell interaction.

|  |  |  |  |
| --- | --- | --- | --- |
|  | Influence on force<br>(attractive or repulsive) |  | Figures |
| *Cellular factors* |  |  |  |
| cell–cell adhesion | attractive |  | 3A |
| cell surface tension | repulsive |  | 3B |
| area elasticity | neutral | core stiffness<br>increased? | 3C |
| cell growth | repulsive |  | 6 |
| *Non-cellular factors* |  |  |  |
| cell-ECM adhesion | repulsive |  | 4 |
| cell-ECM traction force |  |  |  |
| i. random direction | neutral/repulsive |  | 5A |
| ii. uniform | neutral |  | 5B |
| egg-shell | neutral |  | Ref. 1 |
| cavity |  |  |  |
| i. expanding | repulsive |  | Ref. 2 |
| ii. non-expanding | neutral |  | Ref. 2 |

Ref. 1; (Koyama et al., 2023). Ref .2 ; (Koyama et al., 2024)

On the other hand, under the steady states, the relationship between the effective attractive/repulsive forces and the cell–cell adhesion forces/the cell surface tensions was changed in the vertex model (Fig. 7). Vertex models assume many-body interactions but not pairwise interactions (Bi et al., 2016): the potential energies are determined through the relationship among >2 cells. We guess that the effect of the many-body interactions become dominant under the steady states.

Our inference method may be useful for model simplification (i.e., model reduction): models with a higher degree of freedom, such as the vertex models, are simplified as the particle model. Because the particle models are useful to simulate three-dimensional systems as described in the Introduction, it is worth addressing whether three-dimensional models based on the vertex models, etc. can be simplified as the particle models (Kajita et al., 2003; Okuda et al., 2015; Runser et al., 2024).

Some of our theoretical findings were consistent with the cases in MDCK cells with high self-migratory activities or under the steady states (Fig. S3 and S5). To experimentally test the effects of the sub-cellular mechanical components, we need to prepare cells expressing different doses of related proteins such as cadherins for the cell– cell adhesive forces, actomyosin for the cell surface tensions, integrins for the cell–ECM adhesive forces, and so on.

Because real tissues have many mechanical components, it is challenging to separately measure or infer each mechanical parameter value. In the case of our inference method, it may be difficult to distinguish between the increase in the cell–cell adhesive forces and the decrease in the cell surface tensions. To overcome this issue, simultaneous experimental analyses are important: e.g., microscopic imaging of the proteins related to the cell–cell adhesions or the cell surface tensions, such as cadherins, actomyosins, etc. This is not only the case for image-based inference methods but also for invasive methods such as laser ablation experiments, AFM, etc. For instance, by performing the laser ablation experiments, cell–cell junction tensions are estimated (Fernandez-Gonzalez et al., 2009; Ishihara and Sugimura, 2012; Koyama et al., 2016); because the cell–cell junction tensions are composed of both the cell–cell adhesive forces and the cell surface tensions, it is difficult to dissect to what extent the two mechanical components contribute to the cell–cell junction tensions. The Young’s modulus of cells can be measured by AFM, which may be affected by cell elasticities including the area elasticities, and by the cell surface tensions, etc. Therefore, it is important to collect information that describes how the various mechanical components relate to each other. To this end, the theoretical analyses presented in our manuscript, together with the measurements in real tissues, will be invaluable. Although many techniques for measuring/inferring mechanical parameters have been developed, it is still challenging to do in real three-dimensional tissues. This is because of both the accessibility to the tissues and three-dimensional imaging with subsequent image processing (i.e., segmentation). We think that our cell position-based method can overcome these difficulties with parallel applications of the other methods.

## 5. Conflict of Interest

The authors declare that the research was conducted in the absence of any commercial or financial relationships that could be construed as a potential conflict of interest.

## 6. Acknowledgement

We thank Ms. Azusa Kato for supporting nuclear tracking, and Drs. Kazuhiro Aoki, Yohei Kondo, and Kei Yamamoto for advices about the MDCK culture on polyacrylamide gel.

## 7. Author contribution

Conceptualization (HK, AI, HO), Data curation (HK), Formal analysis (HK), Funding acquisition (HK, TO), Investigation (HK, AI, HO, TO, KN, TF), Methodology (HK, KN), Project administration (HK), Resources (HK, TO, TF), Supervision (TF), Validation (HK), Visualization (HK), Writing – original draft (HK), Writing – review and editing (HK, AI, HO, TO, KN, TF)

## 8. Funding

This work was supported by following grants: the National Institutes of Natural Sciences (NINS) program for cross-disciplinary science study for H.K., a Japan Society for the Promotion of Science (JSPS) Grant-in-Aid for Young Scientists (B) for H.K. (17K15131), for Scientific Research (C) for T.O. (18K06234), and for Young Scientists (B) for T.O. (16K18544), and for Transformative Research Areas (A) for HK (24H01990, 24H01414), a MEXT/JSPS Grant-in-Aid for Scientific Research on Innovative Areas for T.O. (17H05627), the Inamori Foundation for T.O., and the Takeda Science Foundation for T.O.

## 9. Declaration of generative AI and AI-assisted technologies

During the preparation of this work, the authors used DeepL in order to improve the readability of the manuscript. After using this tool, the authors reviewed and edited the content as needed and take full responsibility for the content of the published article.

## Notes

### Competing Interest Statement

The authors have declared no competing interest.

### Summary of Updates

A few supplementary data have been added. Small modifications in both the figures and manuscript.

