## Supplementary information for "Cell position-based evaluation of mechanical features of cells in multicellular systems"

Koyama et al.

#### Contents


1. **Generation of synthetic data by particle model**

   1-1. Force fluctuation and its persistency

   1-1. Simulations with traction force between cells and ECM

2. **Generation of synthetic data by vertex model**
3. **Generation of synthetic data by 2-particle for 1-cell model**
4. **Data analysis; definition of heat map**
5. **Experimental procedures for MDCK cells**

   5-1. MDCK strain
   5-2. MDCK culture
   5-3. Acquisition of nuclear tracking data from MDCK cells
   5-4. Immunohistochemistry of ZO-1 and desmoplakin, and staining of cell membrane
   5-5. Measurement of mean square displacement (MSD)

6. **References**
7. **Supplementary Figures**

### 1. Generation of synthetic data by particle model

#### 1-1. Synthetic data from particle model

We performed simulations using distance–force curves. The equation of the particle motion was defined as follows: $V_{C|p} = F_{C|p} / \gamma_C$ (Eq. S1), where $V_{C|p}$ is the velocity of a particle, *p* is an identifier for particles, $F_{C|p}$ is the summation of the cell–cell interaction forces exerted on the *p*th particle, and $\gamma_C$ is the coefficient of the viscous drag and frictional forces. We assumed the simplest situation, i.e., $\gamma_C$ is constant (=1.0) as described previously (Koyama et al., 2023). The simulations were performed by the Euler method written in C.

##### 1-1-1. Force fluctuation and its persistency

We generated simulation data as described previously (Koyama et al., 2023). Briefly, these simulations were performed under the distance–force (DF) curves derived from the Lennard-Jones potential, which can be written as follows; $F(D) = \frac{\kappa}{\sigma_0}\left\{12\left(\frac{\sigma_0}{D}\right)^{13} - 6\left(\frac{\sigma_0}{D}\right)^{7}\right\}$ as a DF curve. $\sigma_0$ corresponds to the distance where the potential becomes 0, and $\kappa$ determines the scale of the forces. In addition, we introduced fluctuations of the cell–cell interaction forces as relative values of the forces derived from the DF curve as follows: $F_{wfl}(D) = F(D) + \nu(pc, \tau)\,|F(D)|$, where $F_{wfl}$ is the force of the cell–cell interactions, wfl means "with fluctuation", and $\nu$ is the magnitude of the fluctuations defined as a ratio to $|F(D)|$. $\nu$ was defined from minus to plus values. $\nu$ is given by Gaussian distribution, $N(0, (pc/100)^2)$, where the average and standard deviation (SD) of the Gaussian distribution is 0 and $(pc/100)$, respectively. *pc* means the percentage. $\tau$ is the persistence time period reflecting time scale of cellular processes. The value of $\nu(pc,\tau)$ is evolved for every time period $\tau$ and is unchanged during $\tau$. In the related figures, *pc* and $\tau$ are shown as "SD value of force fluctuation (%)" and "Persistency of force fluctuation (min)", respectively. The precise definition was described in our previous work (Koyama et al., 2023). $\sigma_0$ was set to 5.0 as the same setting in our previous study (Koyama et al., 2023).

##### 1-1-2. Simulations with traction force between cells and ECM

In Fig. 5, the particles receive the traction forces from the ECM. The movements of the particles were determined by the summation of the forces derived from the DF curves of the cell–cell interactions and of the traction forces. In other words, the traction forces were added to $F_{C|p}$ in the Equation S1. Under the condition that the random vectors of the traction forces were assumed (Fig.

5A), the directions of the vectors were set to be random among the particles, while the magnitudes were set to be the same for all particles. The direction of the vector is evolved for every time period of a given value: the direction is unchanged during the time period. Under the condition that the uniform vector of the traction forces was assumed (Fig. 5B), the traction forces exerted to all particles have the same vector: both the directions and the magnitudes of the vectors were the same for all particles. Thus, the systems move unidirectionally along the vectors, whereas the relative positional relationship among the particles were changed due to the forces from the DF curves.

#### 2. Generation of synthetic data by vertex model

The vertex model used in Fig. 2-3 is essentially the same as that in our previous studies (Koyama et al., 2022; Suzuki et al., 2017). The potential energy in the whole system was defined as: $U = \sum_{<k,l>} \lambda_{\mathrm{Tot}} L^{<k,l>} + \sum_{n} \frac{1}{2} K_{\mathrm{a}} \left( \frac{a_n}{a_{\mathrm{O}}} - 1 \right)^2 a_{\mathrm{O}}$, where each parameter is defined in the main text. In Fig. 4, a following potential was added to the above potential energy: $-\sum_{n} E_{\mathrm{ECM}} a_n$. The force ($F_{\mathrm{V}|h}$) exerted on the $h$th vertex was calculated as follows: $F_{\mathrm{V}|h} = -\nabla U$, where $\nabla$ is the nabla vector differential operator at each vertex. The motion of each vertex is overdamped by the friction forces and is described as follows: $V_{\mathrm{V}|h} = F_{\mathrm{V}|h} / \gamma_{\mathrm{V}}$, $V_{\mathrm{V}|h}$ is the velocity of the $h$th vertex, and $\gamma_{\mathrm{V}}$ is the coefficient of the friction for a vertex. The fluctuation of the tensions ($\sigma$) was provided as follows. The cellular edges to be activated are selected according to $p_{\mathrm{ED}}$ which is the probability of activation per unit of time. On the cellular edges selected, $\sigma$ is added to the line tensions, and the duration time of this activated state is $\tau$.

In the case of Fig. S1, the potential energy based on cell perimeters was assumed: $\sum_{n} \frac{\lambda_{\mathrm{PR}}}{2} \left( L_{\mathrm{PR}}^{<n>} \right)^2$, where $L_{\mathrm{PR}}^{<n>}$ is the perimeter of the $n$th cell, and $\lambda_{\mathrm{PR}}$ is the coefficient. This potential increases the cell–cell junction tensions in a manner dependent to the lengths of the cell perimeters. Parameter values: $\lambda_{\mathrm{S}} = 0.5$ in Fig. 2C, 3A, 3C, 4, and S1B; $\lambda_{\mathrm{AD}} = 0.8$ in Fig. 2C, 3B, 3C, and S1C; $K_{\mathrm{a}} = 1.0$ in Fig. 2C, 3A, 3B, 4, and S1; $\sigma = 0.04$ in Fig. 3-4 and S1 (corresponding to 20%

of $\lambda_{Tot}$ in the case of Fig. 2C, 3C, and 4); $a_o = 25\mu m^2$ in Fig. 2-4 and S1; $p_{ED} = 0.01[min^{-1}]$; $\tau = 20$min in Fig. 2-4 and S1.

#### 3. Generation of synthetic data by 2-particle for 1-cell model

The 2-particle for 1-cell model in Fig. 6 is based on previous studies (Basan et al., 2011; Podewitz et al., 2015) with a few modifications as follows: the velocity of a particle was calculated by the net force exerted on the particle in the same way as Equation S1 with the same value of $\gamma_C$ (=1.0), and the interactions between the particles of different cells were provided by the LJ potential.

#### 4. Data analysis; definition of heat map

The heat maps in Figure 2C were generated from the frequencies of the data points as defined in our previous report (Koyama et al., 2023); the graph space composed of the distance and the force was divided into 64×64 square regions, and then, the data points were allocated to one of the 64×64 square regions. We considered 64 columnar regions along the distance, and calculated the mean values of the counts of the square regions for each columnar region. The mean count for each 64-column region was defined as the frequency index (FI) = 1. The lookup table of the colored map is shown in Figure 2C; many data points were plotted around the red regions, whereas few data points were plotted around the black regions. In the white regions, no data points were plotted. By binned averaging for each column along the distance, DF curves were calculated (Koyama et al., 2023).

#### 5. Experimental procedures for MDCK cells

##### 5-1. MDCK strain

MDCK cells were described previously (Otani et al., 2019), and were originally provided from Masayuki Murata (University of Tokyo). We confirmed the identity of the cell line (derivation from canine) by genomic sequencing of targeted loci. The mycoplasma was removed, and the removal was confirmed by Minerva biolabs Venor GeM OneStep Mycoplasma detection Kit for Conventional PCR.

#### 5-2. MDCK culture

MDCK cells were cultured in DMEM at 37°C. Hoechst 33342 (Molecular probes, Eugene, Oregon, USA) was applied (final conc. = 0.2 μg/mL) to label the nuclei, and the cells were cultured for 1 hr. Then, the medium was replaced with fresh DMEM, and microscopic imaging was performed. Five or six distinct colonies were analyzed for α-catenin mutant or wild-type cells, respectively.

In the case of Fig. S3 or S5, MDCK cells were cultured on glass-bottom dishes (Iwaki, 35mm/ϕ12mm, Japan) or on polyacrylamide gel coated by type I-C collagen (Cellmatrix, Nitta gelatin, Japan). The gel is softer than the glass surface, leading to reduction of the self-migratory activities which are provided from cell–substrate interactions. In Fig. S3, the cells showed a colony, and therefore, they were under the proliferating phases. In Fig. S5, the cells nearly reached confluent states. To reduce self-migratory activities, the serum was removed from the culture medium in Fig. S5.

The polyacrylamide gel was prepared as previously described (Aoki et al., 2017). The gel solution was prepared with 4.0% acrylamide, 0.1% bisacrylamide, 0.08% ammonium persulfate, 0.08% TEMED. 13 ml of the solution was put on a glass bottom dish, and then covered with a glass cover slip with 15mm diameter. After polymerization, the surface was treated with 4mM Sulfo-SANPAH (Sigma) subsequent UV (365nm) irradiation for 10min. Then, after washing out, the surface was coated with the collagen solution (>12 hours).

#### 5-3. Acquisition of nuclear tracking data from MDCK cells

The confocal images obtained from MDCK cells were subjected to the procedure for nuclear detection and tracking. Using the Imaris software (Oxford instruments/Bitplane, UK), the nuclei were automatically detected, followed by manual corrections. The nuclear tracking was also performed automatically, followed by manual corrections. Through the manual corrections, the accuracies of the nuclear detection and tracking became essentially 100%, to the best of our judgment. In the case of the α-catenin mutant MDCK cells, the cells often dissociated from both the dishes and other cells, resulting in free drift. After starting free drift, the cells were considered lost, and were excluded from our analyses.

#### 5-4. Immunohistochemistry of ZO-1 and desmoplakin, and staining of cell membrane

α-catenin knockout (KO) cells were expected to exhibit compromised cell–cell adhesion due to the reduction of cadherin function. The α-catenin mutant cells were constructed previously (Otani et al., 2019). By performing microscopic imaging of cell membrane and of proteins located on cell–cell contacts, we confirmed that the α-catenin KO cells were defective in cell–cell adhesion as follows. Localizations of ZO-1 (a protein located at tight junctions) and localization of desmoplakin (a protein located at desmosomes) were analyzed by performing immunostaining (Fig. S5). These proteins were clearly detected on cell–cell junctions in wild-type cells, whereas these proteins were less or partially detected on cell–cell junctions in the α-catenin KO cells. The cell membranes were stained with FM4-64 fluorescent dye (Fig. S5). In wild-type cells, the cell membranes of neighboring cells were attached each other so that the cell boundaries were shared between the neighboring cells. By contrast, in the α-catenin KO cells, such shared regions were significantly reduced. There results indicate that the cell–cell adhesion in the α-catenin KO cells is compromised.

The protocols of the above experiments are shown as follows. MDCK cells were cultured on glass-bottom dishes (Iwaki, 35mm/ϕ12mm, Japan) for several days. For immunostaining of ZO-1 and desmoplakin, the cells were fixed by methanol for 20 min at -20 °C. After washing with PBS, the cells were permeabilized with 0.1% Triton X-100 for 15 min at room temperature (RT). Blocking was performed using Blocking One (nacalai tesque, Kyoto, Japan) for 10 min at RT. The cells were incubated with the antibodies against ZO-1 (T8-754 (Itoh et al., 1991)) or against desmoplakin (#65146; PROGEN) in 5% Blocking One in PBS for 3 hours at RT. After washing with PBS, the cells were incubated with both a secondary antibody (goat anti-mouse IgG (H+L) Alexa Fluor 594 (A-11032; Invitrogen)) and Hoechst 33258 in 5% Blocking One in PBS for 30-50 min at RT. After washing with PBS, fluorescence images were acquired on a Nikon A1 laser scanning confocal microscope (Nikon, Japan) equipped with a 20× objective (PlanApo; Dry; NA=0.75, Nikon, Japan). For staining by FM4-64, the cells were cultured with both FM$^{TM}$4-64FX (5ng/μL) (F34653; Invitrogen) and Hoechst 33342 for 1 hour, and then, microscopic imaging was performed using the Nikon A1 confocal microscopy. Both the secondary antibody and FM4-64 were excited using a 561-nm laser, and Hoechst 33258 and 33342 were excited using a 403-nm laser. 10 z-slices separated by 0.875μm were imaged, and maximum intensity projection (MIP) images were generated.

#### 5-4. Measurement of mean square displacement (MSD)

α-catenin KO cells migrated on cultured dishes because they have a self-propelled activity originated from their traction forces between the cells and substrates. In comparison with wild-type

MDCK cells, the movements of α-catenin KO cells seemed to be less coordinated, resulting in random walk-like movements. The features of the movements were analyzed by calculating the mean square displacement (MSD). α-catenin KO cells showed increased MSD compared with wild-type cells (Fig. S5C), indicating that α-catenin KO cells move faster than wild-type cells, and are highly diffusive.


#### 7. Supplementary Figures

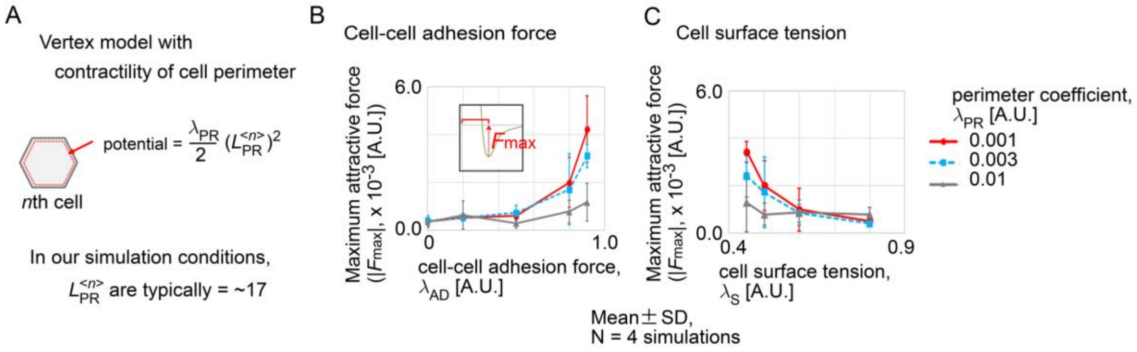


Figure S1 (related to Figure 3): Model dependency of vertex model

A. The potential derived from the cell perimeter is shown, which provides the contractility. $L_{PR}^{<n>}$ is the perimeter of the *n*th cell, and $\lambda_{PR}$ is the coefficient.

B-C. The maximum attractive forces in the inferred DF curves are shown in a manner similar to those in Fig. 3A-iii and 3B-ii, except that $\lambda_{PR}$ was considered. We evaluated the effects of both the cell–cell adhesion forces (B) and the cell surface tensions (C) in this vertex model. Under the smaller values of $\lambda_{PR}$ (red and blue lines), the relationships between the two mechanical components and the attractive/repulsive forces were similar to that in Fig. 3BC, whereas, under the large values of $\lambda_{PR}$ (gray lines), the relationship became ambiguous.

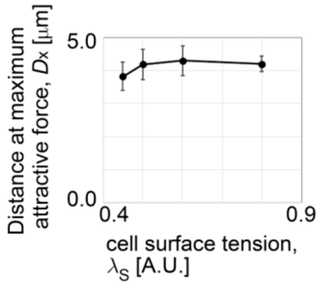


Figure S2 (related to Figure 3B and 4): Evaluation of profile of inferred DF curve

The distances at the maximum attractive forces in the inferred DF curves of Fig. 3B are shown (vs. Fig. 4-iv, the right panel). The cell surface tensions did not significantly affect the distances.

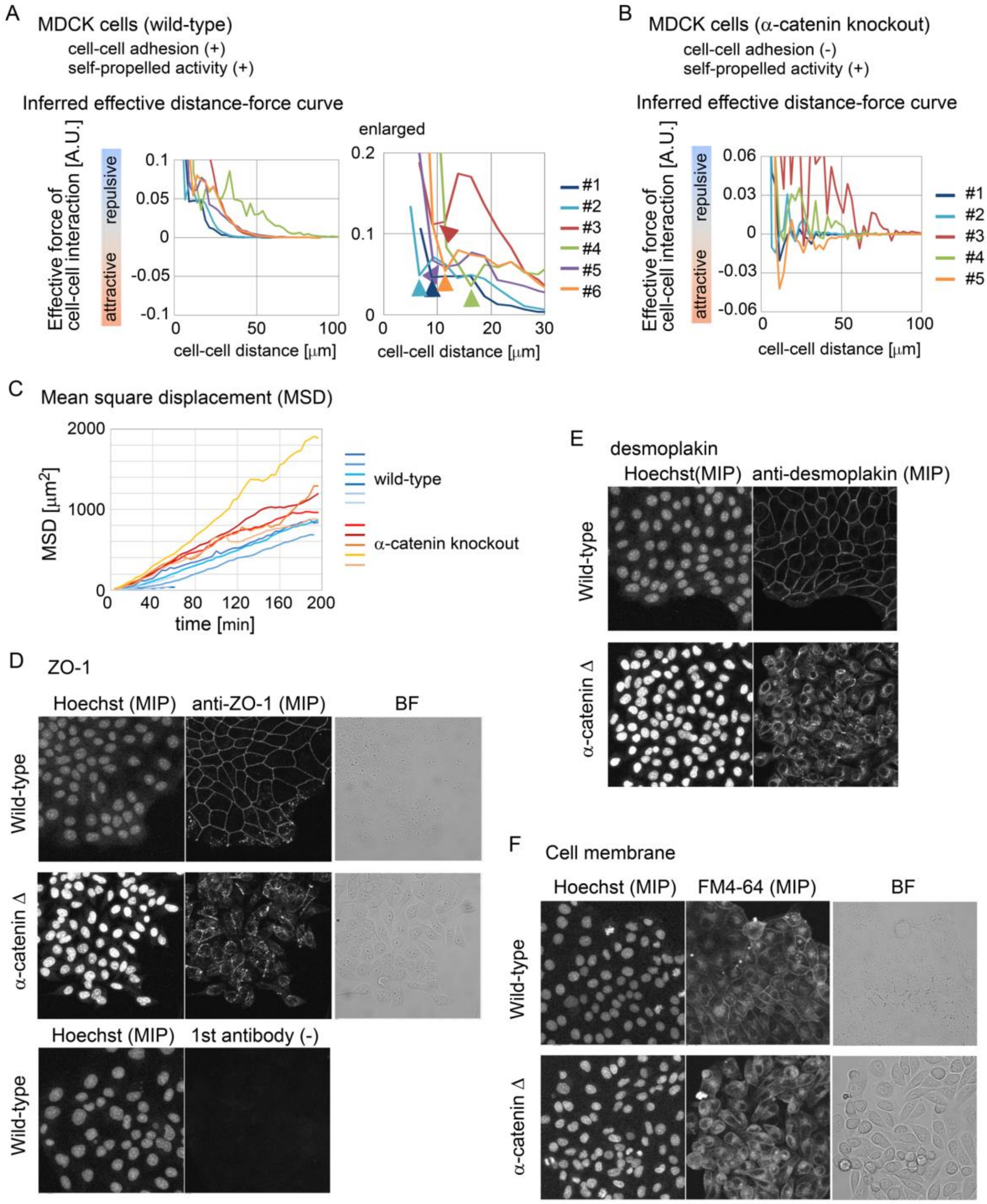


Figure S3 (related to Figure 5): Inference of effective forces in two-dimensionally cultured MDCK cells

A. The effective forces of cell–cell interactions in wild-type MDCK cells are shown, that were two-dimensionally cultured on a dish. The cells are expected to have both the cell–cell adhesion ability and the self-migratory activity. Six independent cell colonies were analyzed. The arrowheads indicate the local minima of the forces, which may be derived from the attractive forces between the cells.

B. The effective forces in α-catenin–knockout MDCK cells are shown, which are expected to have the reduced cell–cell adhesion and to undergo self-propelled movements. Five independent cell colonies were analyzed. The curves are not smooth, resembling the situation in the random walk (Koyama et al., 2023).

C. The mean square displacements (MSD) in wild-type and α-catenin–knockout cells for each colony in A and B are shown. The slopes in the latter cells were steeper than those in the former cells, suggesting that the latter cells diffused faster probably due to the decreased cell–cell adhesion.

D. The confocal images of immunostaining of ZO-1 (a protein localized at tight junctions) in both wild-type and α-catenin–knockout cells (α-cateninΔ) are shown. Hoechst (for nuclear staining); BF, bright field images; MIP, maximum intensity projection image; 1st antibody (-), no anti-ZO-1 antibody as a negative control. The localization of the ZO-1 proteins on the cell–cell junctions was clearly detected in wild-type cells, whereas the localization was reduced or partial in the α-cateninΔ cells.

E. The confocal images of immunostaining of desmoplakin (a protein localized at desmosomes) in both wild-type and α-cateninΔ cells are shown. The localization of the desmoplakin proteins on the cell–cell junctions was clearly detected in wild-type cells, whereas the localization was reduced or partial in the α-cateninΔ cells.

F. The confocal images of the cells stained by FM4-64 (a chemical for staining cell membranes) are shown in both wild-type and α-cateninΔ cells. The cell boundaries detected by FM4-64 were largely shared between the neighboring cells in wild-type cells, whereas a small portion of the cell boundaries were shared in the α-cateninΔ cells. D-F indicate that the cell–cell interactions in the α-cateninΔ cells were severely compromised.

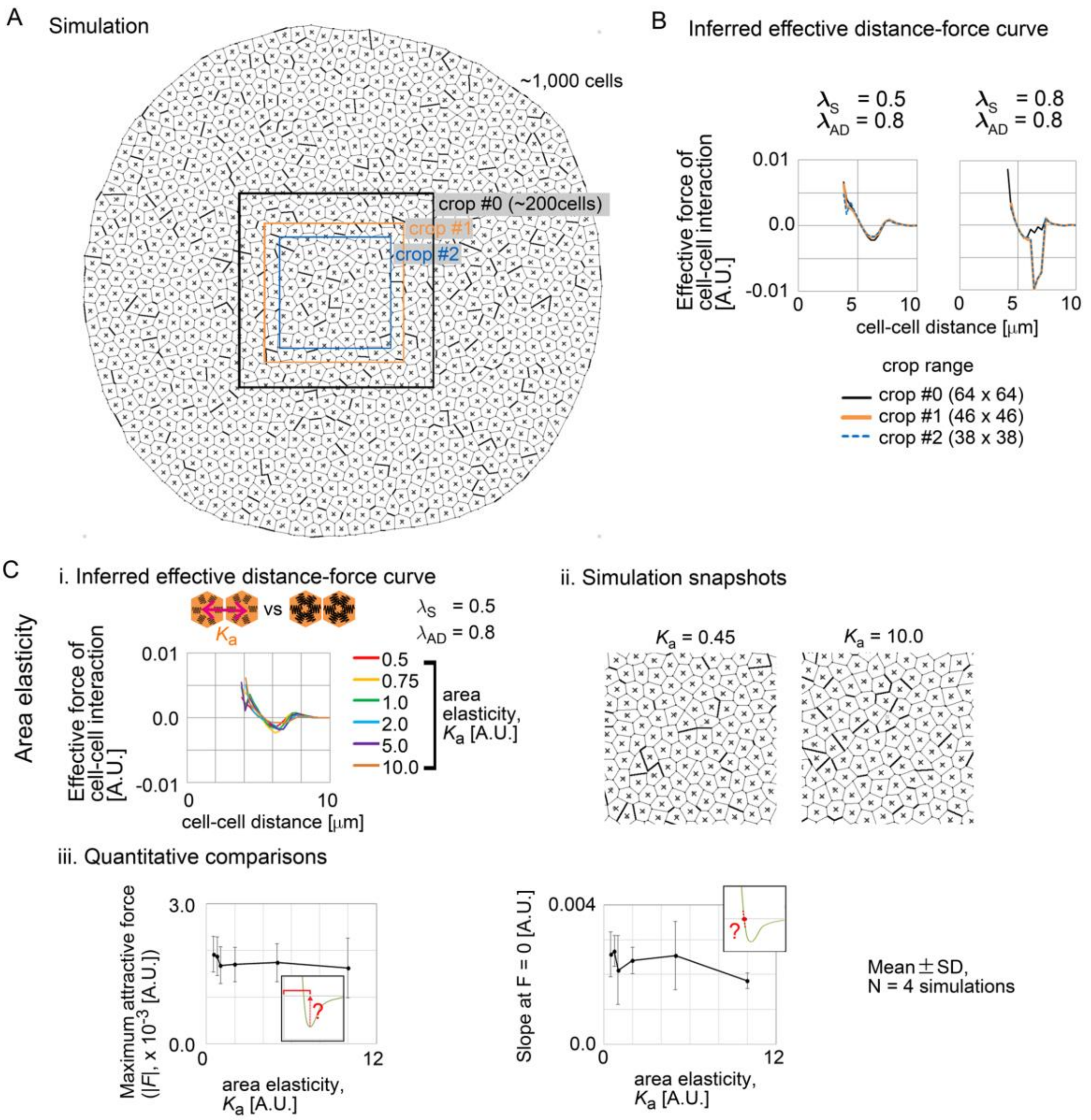


Figure S4 (related to Figure 7): Whole view of simulations and influence of area elasticity on effective attractive/repulsive forces

A. A whole view of a simulation. The effective forces were inferred by using cell tracking data in the "crop #0" region. Then, to reduce the influence of the boundary on the inferred forces, the DF curves were calculated from the effective forces in the smaller inner regions ("crop #1" and "crop #2" regions).

B. Comparison of the inferred DF curves from the crop #0, #1, and #2 regions. The DF curves in the crop #0 can be different from those in the crop #1 and #2 (right panel). But, the DF curves in the crop #1 and #2 were equivalent each other, indicating that the influence of the boundary was almost diminished by cropping smaller regions.

C. Influence of area elasticity. i) Inferred DF curves under different values of the area elasticities. ii) Simulation snapshots. iii) Quantitative comparisons of the profiles of the inferred DF curves.

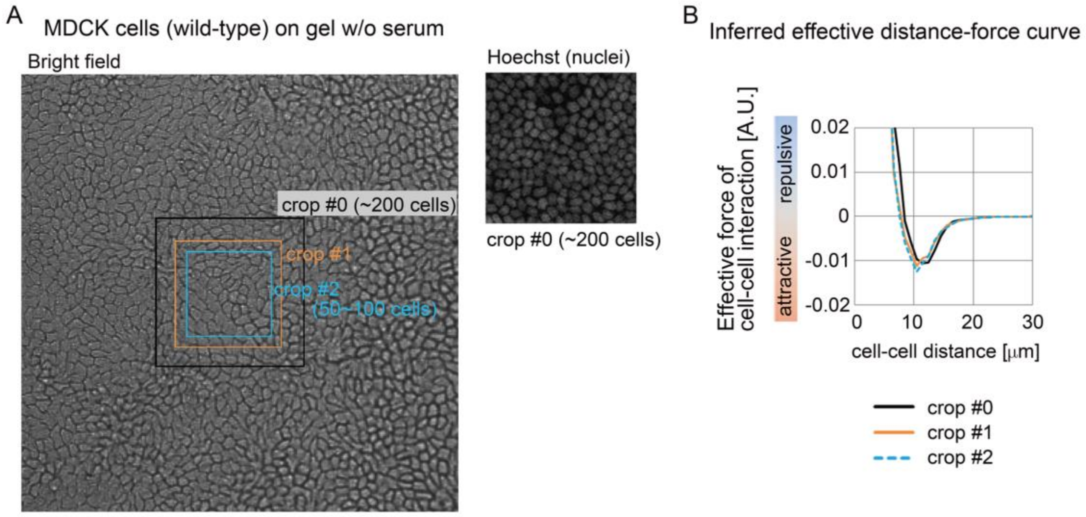


Figure S5 (related to Figure 7): Effective attractive/repulsive forces in MDCK cells under steady state

A. A whole microscopic view of MDCK cells. The cells were cultured on polyacrylamide gel coated by collagen in the absence of the serum. The crop #0, #1, and #2 regions are defined in a manner similar to the case in Fig. S4.

B. Inferred DF curves from the crop #0-2 regions in a manner similar to the case in Fig. S4. The DF curves from the crop #1 and #2 were well consistent each other, whereas the curve from the crop #0 showed a slight difference from those from the crop #1 and #2.
